# The Size-Weight Illusion is unimpaired in individuals with a history of congenital visual deprivation

**DOI:** 10.1101/2021.02.10.430572

**Authors:** Rashi Pant, Maria J. S. Guerreiro, Pia Ley, Davide Bottari, Idris Shareef, Ramesh Kekunnaya, Brigitte Röder

## Abstract

Visual deprivation in childhood can lead to lifelong impairments in visual and multisensory processing. Here, the Size-Weight-Illusion was used to test whether visuo-haptic integration recovers after sight restoration. In Experiment 1, congenital (CC: 7 (3F), 8–35 years) and developmental cataract reversal individuals (DC: 9 (2F), 8–37 years), as well as congenitally blind (CB: 2 (1F), 33 and 44 years) and normally sighted individuals (SC: 10 (7F), 19-36 years) perceived larger objects as lighter than smaller objects of the same weight. In Experiment 2, CC (6 (1F), 17–44.7 years) and SC (7 (5F), 21-29 years) individuals performed identically when tested without haptic size cues. Together, this suggested that early visual experience is not necessary to perceive the Size-Weight-Illusion.

---

Infants born with dense bilateral cataracts lack patterned vision until their sight is restored by cataract removal surgery. When surgery is performed late, i.e. beyond the first few weeks from birth, these individuals have been reported to suffer from permanent visual and multisensory impairments (Birch, Stager, Leffler, & Weakley, 1998; de Heering et al., 2016; Lewis & Maurer, 2005; Lewkowicz & Röder, 2015; Maurer, 2017). These impairments are hypothesized to be a behavioral consequence of neural system changes, resulting from aberrant sensory input within a sensitive period of development (Knudsen, 2004; Maurer, 2017). One way to assess visual and multisensory functional recovery in cataract reversal individuals is to test their susceptibility to well-known perceptual illusions. Perceptual illusions are typically extremely robust, suggesting that they arise from automatic processing principles (Eagleman, 2001). Thus, the lack (or reduced likelihood) of perceiving a visual or multisensory illusion is indicative of impaired visual or multisensory processing, respectively.

Putzar et al. (2007) tested the automatic detection of illusory contours in congenital cataract reversal individuals who had undergone cataract surgery between 1 and 17 months of age (Putzar et al., 2007). They observed deficits in individuals who experienced more than 5-6 months of visual deprivation, and interpreted this result as evidence for impairments in the automatic binding of visual features – i.e. visual feature binding. This finding was extended by McKyton et al (2015) to a population of cataract reversal individuals who had undergone surgery only after the age of 5 years (McKyton et al., 2015). These results suggested that early visual experience is required to acquire the neural circuits necessary for visual feature binding, providing evidence in favor of the sensitive period hypothesis. While this conclusion is compatible with the long developmental trajectory of illusory contour perception (Hadad, Maurer, & Lewis, 2010), other studies replicated this finding for individuals treated for monocular (but not binocular) congenital cataracts (Hadad, Maurer, & Lewis, 2017).

Gandhi et al. (2015) tested the Ponzo and Müller-Lyer illusions, wherein equally long lines are perceived to be of different lengths depending on their surroundings (Gandhi, Kalia, Ganesh, & Sinha, 2015). Forty-eight hours after cataract surgery, they found a full recovery of this illusion, despite the fact that sight was partially restored only at 8 years of age or later (Gandhi et al., 2015). Since both these illusions were thought to arise from the interpretation of two-dimensional perspective cues as three-dimensional depth, they concluded that this process did not depend on early childhood vision.

Illusions have additionally been used to investigate the extent of multisensory recovery following cataract surgery. In an initial study, Putzar et al. (2007) employed an audio-visual temporal capture effect, characterized as follows: if an auditory stimulus is presented with a short temporal offset with respect to a visual stimulus, the visual stimulus is often perceived as temporally shifted towards the time point when the sound was presented (Putzar, Goerendt, Lange, Rösler, & Röder, 2007). This effect was significantly reduced in congenital cataract reversal individuals (Putzar, Goerendt, et al., 2007). The residual visual impairments would have, according to the inverse efficiency rule of multisensory integration, predicted a larger capture in this group, suggesting that the multisensory binding process was impaired (Meredith & Stein, 1983). Similarly, cataract reversal individuals had a significantly reduced likelihood of showing the McGurk effect (Putzar, Hötting, & Röder, 2010). In this illusion, auditory speech presented concurrently with an incongruent visual lip movement produces a percept that matches neither the auditory nor the visual input. The absence of the McGurk effect in congenital cataract reversal individuals was subsequently shown to be related to a lack of multisensory enhancement in superior temporal brain regions known to be essential for audio-visual speech perception (Kumar et al., 2016; Nath & Beauchamp, 2012; Sams et al., 1991). Finally, visual motion after-effects (perception of stationary stimuli as moving in the direction opposite to previously presented moving stimuli) were found after auditory motion adaptation in cataract-reversal individuals, i.e., a cross-modal after-effect which has not been found in normally sighted individuals (Guerreiro, Putzar, & Röder, 2016). These results converge to the conclusion that multisensory binding processes do not fully recover after sight restoration in individuals with a history of congenital cataracts. Although recent studies with cataract reversal individuals have demonstrated recovery of multisensory redundancy effects (de Heering et al., 2016; Putzar, Gondan, & Röder, 2012) and partially for auditory-visual simultaneity judgements (Chen, Lewis, Shore, & Maurer, 2017), multisensory binding based on more complex features, such as speech (Putzar, Goerendt, et al., 2007), seems to depend on early visual input.

It is currently unclear to what degree visuo-haptic and visuo-motor processing recovers after a transient phase of congenital visual deprivation. A developmental study with children found that visuo-haptic integration reaches adult-like performance only by 10 years of age (Gori, Del Viva, Sandini, & Burr, 2008). Prior to that, children show signs of either vision or touch dominating visuo-tactile perception. Evidence from studies in a small number of individuals who had dense (but not necessarily congenital) bilateral cataracts suggested a quick emergence of visuo-haptic interactions after surgery, when these individuals were tested in vision-to-touch object matching tasks (n=1, Chen et al., 2016; n=5, Held et al., 2011). However, later single case studies pointed towards impaired spatial representations for visuo-tactile localization after at least two years of visual deprivation (Azañón, Camacho, Morales, & Longo, 2018; Ley, Bottari, Shenoy, Kekunnaya, & Röder, 2013). Further, a study testing sight recovery individuals on an automatic imitation task, which mapped vision to motoric performance, found performance deficits even two years after surgery (McKyton, Ben-Zion, & Zohary, 2018).

A phenomenon observed in typical visuo-haptic development is the perception of the size-weight illusion (henceforth referred to as the SWI), which has been described as “immutable”(Murray, Ellis, Bandomir, & Ross, 1999). The SWI is an illusion perceived when two unequally sized objects of the same weight are compared - the smaller object is perceived as being heavier than the larger one.

The “classical”SWI is assumed to require the integration of visual and haptic input (for review, see Dijker, 2014). Though the SWI has been documented for a long time, there is still a debate of whether it occurs due to conflicting sensorimotor input, or is a purely cognitive effect due to a mismatch in expectations (Flanagan, Bittner, & Johansson, 2008). At present, the contribution of early visual experience to the occurrence of the illusion is unclear.

Several lines of evidence have suggested a crucial role of ongoing visual input in the perception of the SWI. First, the SWI was reported to disappear when visual cues were not presented to normally sighted participants, even if they were allowed to access them beforehand, suggesting a crucial role of continued visual perception for the illusion, and providing a strong argument that the SWI reflects visuo-haptic integration (Masin & Crestoni, 1988). Additionally, the SWI was observed to increase with an increase in visual disparity between sizes; if two objects of the same weight had a greater difference between their visually perceived sizes, the illusion perceived was stronger (Kawai, Henigman, MacKenzie, Kuang, & Faust, 2007). This was found to be true even in the absence of haptically perceived size differences, when visually perceived size was varied using objects with adjustable heights but constant surface area (Plaisier & Smeets, 2015). Finally, in a rapid adaptation study using functional MRI with an SWI task, the ventral premotor area (PMv) responded more when the SWI was perceived, i.e. when participants compared the weights of two objects of different sizes and the same weight, than when participants compared objects of the same size and weight (Chouinard, Large, Chang, & Goodale, 2009). Importantly, the PMv did not show an independent adaptation to size and weight properties, but adapted to the combination of these properties, therefore providing neural evidence for the integration of concurrent but separate sensory input. These results suggest that early visual deprivation might affect the SWI, if the ability to integrate visual and haptic cues does not recover.

The SWI was observed to occur when size information was perceived exclusively visually (i.e. using a string set-up to weigh objects, preventing individuals from using haptic cues to estimate size) or exclusively haptically (i.e. blindfolding individuals to prevent them from accessing visual estimates of size) (Ellis & Lederman, 1993; Pick & Pick, 1967). However, both these studies reported that the SWI was smaller when the task presented sighted individuals with size cues that were exclusively visual, as compared to when the task was haptic or visuo-haptic. Moreover, congenitally blind individuals have been reported to experience the SWI as well (Ellis & Lederman, 1993; Rice, 1898). These results would lead us to hypothesize that the SWI emerges independent of visual input and thus, should manifest unimpaired in sight-recovery individuals too - at least if the objects are perceived exclusively haptically. However, there might be some competition between the belatedly available visual input and other sensory modalities (Guerreiro, Putzar, & Röder, 2015; Singh, Phillips, Merabet, & Sinha, 2018) which might worsen visual performance (Guerreiro, Putzar, & Röder, 2016; Putzar, Goerendt, et al., 2010). Consistent with this assumption, a reduced visuo-haptic SWI was reported in visually impaired individuals with some residual visual capabilities (Furth, 1961). Therefore, it could be hypothesized that the visuo-haptic SWI, both with and without haptic size information, is reduced in individuals who recover from transient congenital or developmental visual deprivation.

The second account for the occurrence of the SWI has proposed that this illusion is a result of violated expectations, and originates from top-down rather than bottom up processes (Buckingham & Goodale, 2010; Chouinard et al., 2019; Peters, Balzer, & Shams, 2015; Ross, 1966). If the SWI is a result of exclusively visual statistics gathered over the lifespan – i.e. an increased amount of force on the larger object because of the belief that smaller objects should weigh less – then individuals who have not had access to vision early in life, when tested with and without haptic size information, should either not manifest the illusion, or perceive it to a lower degree.

In order to test for the dependence of visuo-haptic interactions on early visual input, we performed two experiments using an SWI paradigm employed by Buckingham and Goodale (2010), wherein subjects were asked to rate cubes on how heavy they thought they were (Buckingham & Goodale, 2010). The first experiment tested the “classic”visuo-haptic SWI, that is the SWI when simultaneously receiving visual and haptic size information. A group of dense bilateral congenital cataract reversal individuals (CC) as well as a group of individuals who had suffered developmental cataracts (DC) were tested, and compared to a group of normally sighted controls (SC). We additionally ran and separately analyzed data from two congenitally blind (CB) individuals to pilot whether we would replicate the findings from Ellis and Lederman (1993). The second experiment used a string set-up to ensure only visual size cues were available to participants, in order to isolate the visual contribution to the SWI (Buckingham, Milne, Byrne, & Goodale, 2015; Ellis & Lederman, 1993; Pick & Pick, 1967). A group of congenital cataract reversal individuals (CC) and a group of normally sighted controls (SC) were compared. We hypothesized that in Experiment 1, the SWI would be impaired in CC compared to SC and DC individuals, due to aberrant visual experience interfering with multisensory integration (Guerreiro et al., 2015; Putzar, Goerendt, et al., 2007), and that in Experiment 2, CC individuals would not experience the SWI when deprived of haptic size information (Buckingham et al., 2015; Ellis & Lederman, 1993).

## Methods

### Ethical Approval

All participants, as well as their legal guardians in case of minors, provided written and informed consent. Testing was conducted after obtaining ethical approval from the German Psychological Society (DGP) and the local ethics board of the Hyderabad Eye Research Foundation. All methods and tests were performed in accordance with the relevant guidelines and regulations of both collaborating institutions.

### Experiment 1

#### Participants

We tested three groups of individuals. The first group consisted of seven individuals born with dense bilateral cataracts, who subsequently underwent cataract removal surgery (referred to as CC group: 3 females, 4 males; Age = 8 – 35 years, M = 21.2 years, SD = 9.8, Table 1). The CC individuals were diagnosed by ophthalmologists and optometrists at the LV Prasad Eye Institute (LVPEI) in Hyderabad (India). They were tested by the some of the authors, partially with the help of a translator, in English, Hindi or Telugu. The data from four additional CC individuals were excluded due to unwillingness to cooperate (n = 2) or a documented developmental delay (n = 2). Individuals were categorized as part of this group based on the presence of dense bilateral cataracts at birth, a pre-surgery visual acuity of counting fingers at 1m or less (barring absorption of lenses), presence of nystagmus, occlusion of fundus/retina, and immediate family members who had also been diagnosed with dense bilateral congenital cataracts. Duration of blindness was calculated by subtracting the date of birth from the date of the first eye surgery (M = 13.21 years, SD = 8.24, Range = 2 – 23.05 years). One individual did not have his precise date of surgery information available (operated after 6 months of age), and was excluded from duration calculations. Visual acuity pre-surgery in the better eye ranged from a minimum of light perception (PL+) to a maximum of counting fingers close to the face (CFCF). One participant had been able to count fingers at a distance of 3m pre-surgery. We included this participant due to clearly partially absorbed lenses (OD Visual Acuity: counting fingers at 1.5m, OS Visual Acuity: counting fingers at 3m). All other criteria such as nystagmus and family history pointed towards the presence of dense bilateral cataracts at birth. Visual acuity post-surgery in the better eye in this group ranged from a minimum of counting fingers at a distance of 1 m to a maximum of 20/40. All individuals included in this group lacked patterned vision at birth, in accordance with the criteria set by the WHO (World Health Organisation, 2019).

**Table 1:** Participant characteristics for all sight recovery participants in Experiments 1 and 2. Age was calculated on the day of testing, and duration of blindness was calculated by subtracting date of birth from date of first surgery. Time since surgery was calculated by subtracting date of first surgery from date of testing. Visual acuity is reported separately for each eye (CF: Counting Fingers, PL: Perception of light, PR: Projection of Rays in all quadrants, FL: Fixate and Follow Light).

| SUB | AGE<br>(YEARS) | GROUP | GENDER | ABSORBED<br>LENSES | PRESENCE<br>OF<br>NYSTAGMUS | PRE-SURGERY VISUAL |  | POST-SURGERY VISUAL |  | DURATION<br>OF<br>BLINDNESS<br>(YEARS) | TIME<br>SINCE<br>SURGERY<br>(YEARS) |
| --- | --- | --- | --- | --- | --- | --- | --- | --- | --- | --- | --- |
|  |  |  |  |  |  | ACUITY |  | ACUITY |  |  |  |
|  |  |  |  |  |  | RIGHT | LEFT | RIGHT | LEFT |  |  |
| EXPERIMENT 1 |  |  |  |  |  |  |  |  |  |  |  |
| 1 | 23.25 | CC | Male | No | Yes | Unknown | Unknown | CF (close<br>to face) | 20/126 | Unknown |  |
| 2 | 35.54 | CC | Male | No | Yes | Unknown | Unknown | 20/120 | 20/40 | 2.00 | 33.54 |
| 3 | 21.78 | CC | Female | Yes | Yes | Unknown | Unknown | 20/317 | 20/252 | 21.02 | 0.76 |
| 4 | 16.97 | CC | Female | No | Yes | PL+ PR+ | CF (Close<br>to Face) | PL+ PR+ | CF at 1m | 16.05 | 0.92 |
| 5 | 30.76 | CC | Male | No | Yes | Unknown | Unknown | 20/400 | 20/800 | 23.05 | 7.72 |
| 6 | 11.24 | CC | Female | No | Yes | PL+ | PL+ | 20/125 | CF at 0.5m | 10.16 | 1.08 |
| 7 | 8.62 | CC | Male | Yes | Yes | CF at 1.5 m | CF at 3 m | 20/500 | 20/126 | 7.00 | 1.62 |
| 8 | 18.34 | DC | Female |  | No | FL+ | FL+ | 20/60 | 20/30 | -- | -- |
| 9 | 37.02 | DC | Male |  | Yes | 20/200 | 20/300 | 20/200 | 20/200 | -- | -- |
| 10 | 10.05 | DC | Male |  | Yes | PL+ PR<br>inaccurate | PL+ PR<br>inaccurate | 20/1200 | 20/1200 | -- | -- |
| 11 | 8.62 | DC | Male |  | No | 20/100 | 20/100 | 20/20 | 20/20 | -- | -- |
| 12 | 8.01 | DC | Male |  | No | 20/80 | 20/100 | 20/40 | 20/40 | -- | -- |
| 13 | 10.15 | DC | Female |  | No | PL+ PR+ | CF at 2m | 20/170 | 20/16 | -- | -- |
| 14 | 17.71 | DC | Male |  | No | 6/60 | CF at 1m | 20/63 | CF at 0.5m | -- | -- |
| 15 | 14.24 | DC | Male |  | Yes | FL+ | Poor<br>fixation | 20/40 | CF at 0.5m | -- | -- |
| EXPERIMENT 2 |  |  |  |  |  |  |  |  |  |  |  |
| 1 | 33.09 | CC | Male | No | Yes | Unknown | Unknown | CF at 1m | 20/200 | 14.00 | 19.10 |
| 2 | 15.24 | CC | Male | Yes | Yes | CF at 1.5m | CF at 3m | 20/500 | 20/126 | 7.00 | 8.24 |
| 3 | 18.30 | CC | Male | Yes | Yes | 20/500 | 20/800 | 20/125 | 20/500 | 16.45 | 1.85 |
| 4 | 17.28 | CC | Female | Yes | Yes | 20/600 | CF at 1m | 20/250 | 20/500 | 15.42 | 1.85 |
| 5 | 37.39 | CC | Male | No | Yes | Unknown | Unknown | 20/400 | 20/800 | 23.05 | 14.34 |
| 6 | 44.74 | CC | Male | Yes | Yes | 20/300 | Unknown | 20/125 | 20/200 | 22.02 | 22.72 |

The second group consisted of nine individuals who had either partial congenital cataracts or developmental cataracts, and were subsequently operated upon to remove the cataracts (referred to as DC group: 2 females, 7 males; Age = 8 – 37 years, M = 14.8 years, SD = 9.2, Table 1) The testing procedure was the same as that of the CC group. Comparing this group with the CC individuals allowed us to isolate effects caused by transient patterned visual deprivation from birth, from effects due to general visual impairments caused by a changed periphery. Visual acuity pre-surgery in the better eye ranged from following light to a maximum of 20/80. Visual acuity post-surgery in the better eye ranged from 20/1200 to 20/20. All individuals included in this group did not lack patterned vision at birth, but suffered from degraded visual input for some or all of their early childhood, therefore providing a control group for the possibility that any observed impairments of the CC group were not specific to visual input at birth, but due to degraded visual input at any stage of life.

The third group consisted of 10 individuals with normal or corrected-to-normal vision and who had no history of visual deficits or eye injuries (referred to as SC group: 7 females 3 males; Age = 19-36 years, M = 25.8 years, SD = 5.3). SC individuals were tested at the University of Hamburg, Hamburg, Germany, in German.

In addition to these three groups, we ran two congenitally blind individuals who had no more than light perception since birth and at the time of the study (referred to as CB group: 1 female, 1 male; Ages = 33 and 44 years). They were tested at the University of Hamburg, Germany, using German. Their data were analyzed separately due to the small group size. The purpose of including these two participants was to replicate the presence of the SWI in individuals who totally lack vision since birth, but not for statistical comparisons between groups (Ellis and Lederman 1993, Buckingham et al 2015). Such an illusion would necessarily be based on exclusively haptic estimates of size.

All individuals included in the data analysis reported no history of neurological or cognitive impairments. All participants were right handed, and used that hand to perform the task.

#### Stimuli and Apparatus

Participants were tested using a free-rating, absolute-magnitude-estimation procedure: they freely chose a rating scale of their preference and estimated how much an object weighed on that scale (Buckingham & Goodale, 2010). This scale was adjustable during the course of the experiment.

Participants rated one of 6 gray plastic cubes that were placed on their palms. The cubes were small (5 cm^3^), medium (7.5 cm^3^) or large (10 cm^3^), and had one of two different weights (either 350g or 700g) (Figure 1). The weight was invisibly fixed in one corner, with the rest of the inside being hollow to ensure the same distribution of weight in each cube. Participants were instructed to hold the dominant arm bent at 90 degrees, and for each trial, the cube was placed on the palm of their dominant hand such that it was clearly visible. Participants were required to lift each cube for approximately 15 seconds, and to judge its weight. Upon the rating response, the experimenter removed the cube from the participants’ hand.

**Figure 1:**
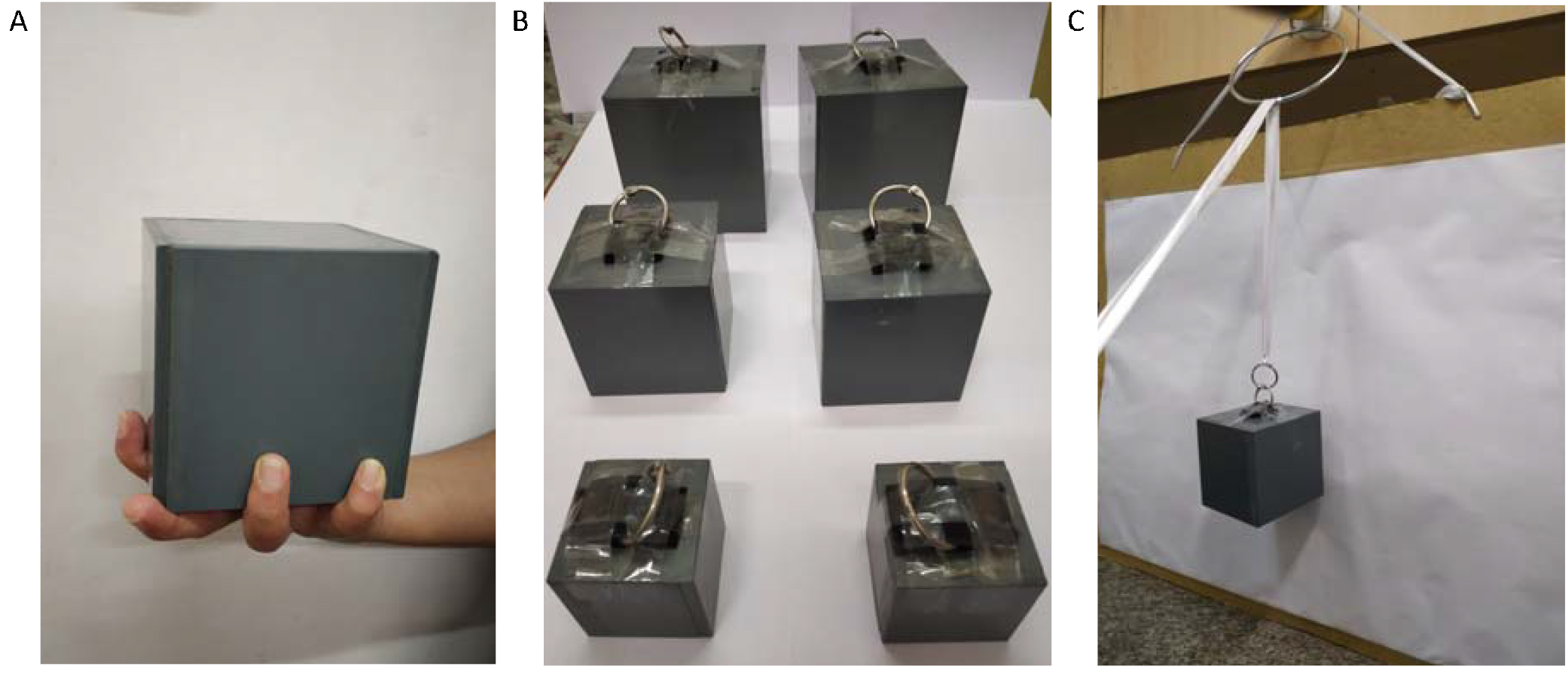
A: Large-sized cube stimulus placed in the dominant hand, demonstrating the procedure for Experiment 1. B: The six stimuli used in the study (front to back: Small, Medium, Large, 350g and 700g respectively), with attachments for Experiment 2. C: String set-up of Experiment 2. A smooth ribbon is used to lift the cube during each trial, ensuring that only visual size cues are available to the participant.

We used random orders, and in one run, each cube was lifted 5 times. We repeated this across two runs, leading to a total of 60 trials. For participants who did not complete 60 trials, we used only the completed run of 30 trials (CC: n=1; DC: n=3). Participants were instructed not to rotate the arm for additional sensory cues, and not to throw the cube in the air and catch it again. This procedure took around 30 minutes in total.

#### Data Analysis

In order to compare subjective judgements across participants, rating scores were z-transformed within each participant. This was done by subtracting the individual’s mean weight judgement from each weight judgement, and dividing by the standard deviation (Buckingham & Goodale, 2010). Due to this z-scoring, all weight judgements reflect deviations from the same mean (zero), with higher z values indicating heavier weight judgements. In order to be included in the data analysis, participants had to consistently rate the 350g cubes as lighter than the 700g cubes, to exclude the possibility of a response bias as opposed to a principled difference in perceived weight due to size. This was true of all participants in Experiment 1.

We used frequentist statistics to analyze the data. Z-scores across participants were submitted to a mixed ANOVA. Our model considered two within-group factors – namely, weight (2 levels: 350g, 700g) and size (3 levels: small, medium, large), and one between-group factor – group (3 levels: CC, DC, and SC) in a repeated measures ANOVA. Levene’s test for Homogeneity of Variance was performed on the z-score data to ensure that the scores do not violate the assumption of equal variance for parametric testing, due to unequal sample sizes between groups (*F(*2,23)=0.27, *p*=0.764). Post-hoc ANOVAs and t-tests were performed according to the resulting interactions, and post-hoc equivalence testing was conducted to confirm the results (Supplementary Information S1).

All analyses were conducted in R (version 3.3.2), using the ez-package (https://github.com/mike-lawrence/ez). This package corrects for violations of sphericity when there are more 2 levels in the within subject variable (size) via the Greenhouse-Geisser correction. All effect sizes reported are generalized eta squared (η_g_^2^) values.

This study was not pre-registered, and sample sizes were limited by strict inclusion criteria within a special population.

## Results

### Sight-recovery individuals show an intact visuo-haptic SWI (Experiment 1)

When z-scored weight ratings were assessed in a size-by-weight-by-group analysis of variance (ANOVA with repeated measures), all groups performed the task in a principled manner, as indicated by a main effect of weight, wherein the 700g weight was rated as heavier than the 350g weight (F(1,23)=639.15, p<0.001, η_g_^2^=0.85). There was a main effect of size, demonstrating the presence of the SWI (F(2,46)=147.11, p<0.001, η_g_^2^=0.74), that is, participants perceived smaller sized objects of the same weight to be heavier. Crucially, the group-by-size (*F*(4,46)= 0.96, *p*=0.419, η_g_ ^2^=0.03), group-by-weight (*F*(2,23)= 2.63, *p*=0.094, η_g_ ^2^=0.04), and size-by-weight-by-group interactions (*F*(4,46)= 1.64, *p*=0.181, η_g_ ^2^=0.05) were all not significant. Therefore, CC individuals displayed an indistinguishable SWI from DC and SC individuals (Figure 2, Figure 3).

**Figure 2:**
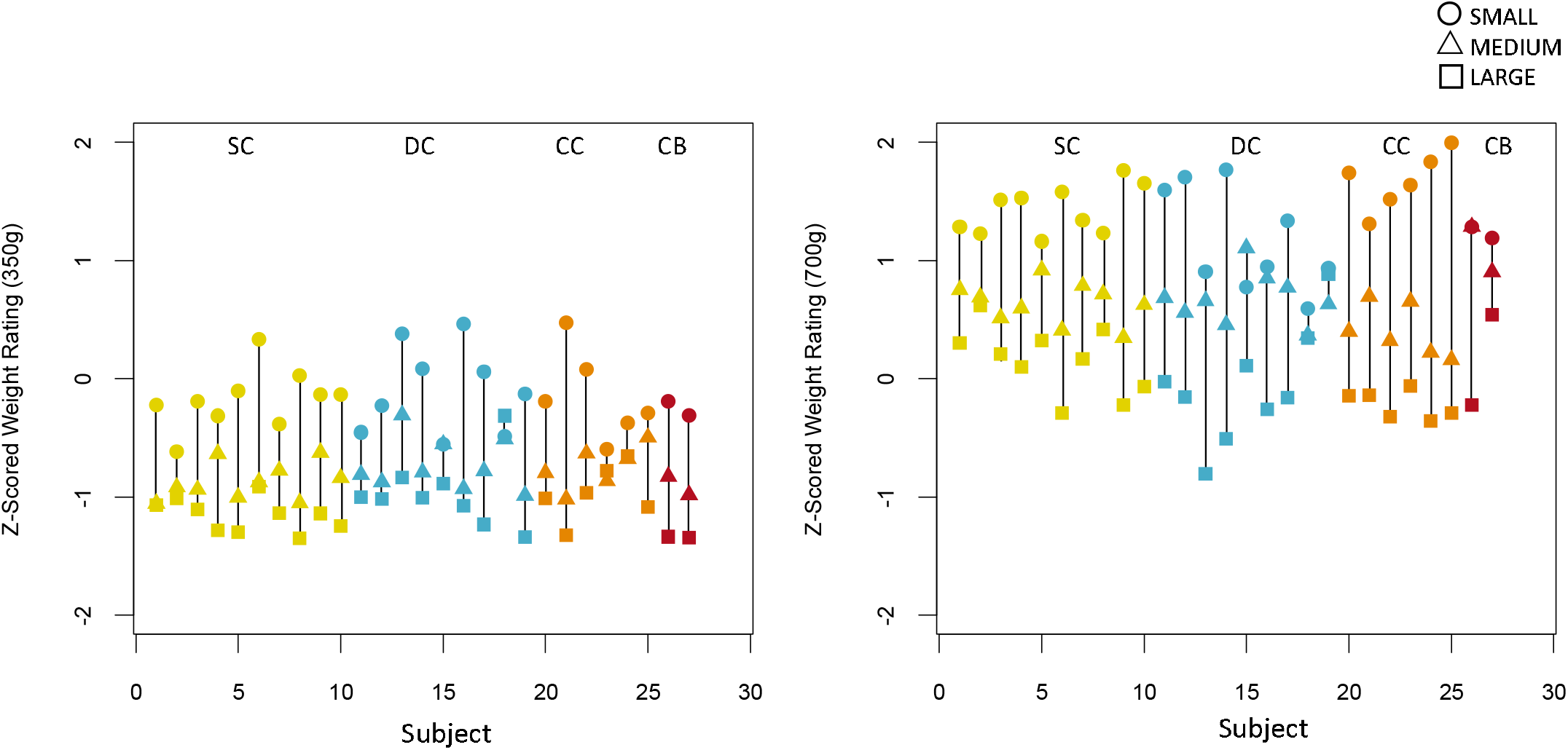
Experiment 1 (with visual and haptic size information); z-scored weight ratings of all individuals in the Sighted Control (SC, yellow), the Developmental Cataract group (DC, blue), the Congenital Cataract group (CC, orange), and two Congenitally Blind (CB, red) individuals, t (A) for the 350g and (B) for 700g weights (Experiment 1).

**Figure 3:**
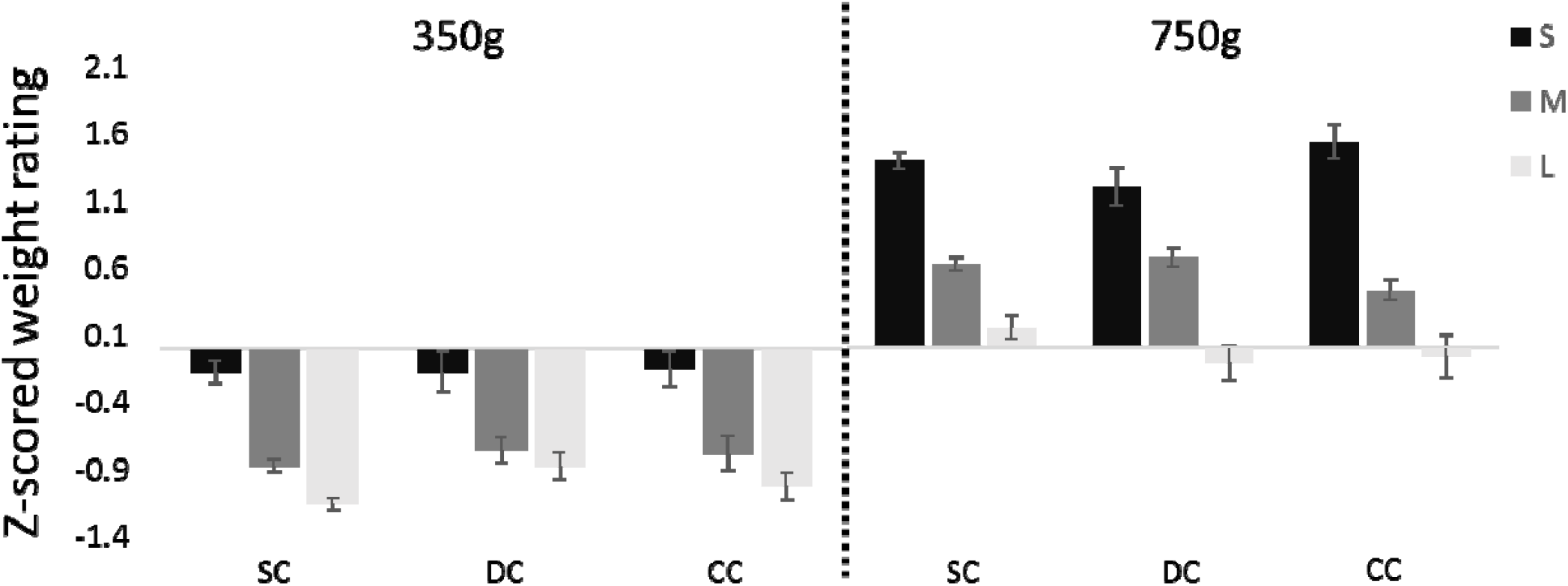
Experiment 1 (with visual and haptic size information); average z-scored weight ratings for the Small (S), Medium (M) and Large (L) cubes across the Sighted Control (SC), Developmental Cataract (DC) and Congenital Cataract (CC) groups, for the 350g and 700g weights (Experiment 1). Error bars depict standard error of mean (SEM).

We found a size-by-weight interaction, indicating that the illusion was stronger for the 700g than the 350g weight (*F*(2,46)= 11.82, *p*=0.001, η_g_ ^2^=0.15). In order to follow up on this effect, two post-hoc 3 (size) X 3 (group) repeated measures ANOVAs were conducted, separated by the weight condition. There was a main effect of size for both the 350g (*F*(2,46)= 59.08, *p*<0.001, η_g_ ^2^=0.66) and the 700g (*F(*2,46)=108.24, *p*<0.001, η_g_ ^2^=0.79) weights, demonstrating that the SWI was highly significant for both weights in all groups. Again, we found no group-by-size interaction (350g: *F*(4,46)=0.57, *p*=0.63, η_g_ ^2^=0.04; 700g: *F*(4,46)=1.74, *p*=0.179, η_g_ ^2^=0.11), indicating that the degree to which the three groups experienced the SWI was indistinguishable.

Paired t-tests across the three groups confirmed that participants experienced the smaller cube as heavier than the medium (350g: *t*(24) = 8.28, *p* < 0.001; 700g: *t*(24) = 7.64, *p* < 0.001) and large cube (350g: *t*(24) = 9.95, *p* < 0.001; 700g: *t*(24) = 11.52, *p* < 0.001) of the same weight, and the medium sized cube as heavier than the large cube of the same weight (350g: *t*(24) = 5.97, *p* < 0.001; 700g: *t*(24) = 8.49, *p* < 0.001).

Both CB individuals tested showed an impressively clear SWI (Figure 2). We found their z-scores (Figure 2) to fall within the core range of the remaining groups. Thus, these two participants replicated the results of Ellis and Lederman (1993).

#### 1.1 SWI Index

In order to obtain a measure of illusion strength for each individual, we calculated an SWI Index by subtracting the mean z-scored weight judgment of the largest cube from that of the smallest cube, separately for each weight.

We conducted separate one way ANOVAs for the 350g and 700g weights to assess the effect of group on the calculated SWI Index. The CC, DC, and SC groups did not differ in the strength of the illusion - neither for the 350g (*F*(2,23)=0.72, *p*=0.497, η_g_ ^2^=0.06) nor the 700g weights (*F*(2,23)=0.74, *p*=0.487, η_g_ ^2^=0.06) (Figure 6). We further tested these null findings against effect sizes obtained from studies in the literature. Based on a comparable study and task found for the 700g weight, we used equivalence analyses to confirm that the SWI index was equivalent in the CC, DC and SC groups – i.e. the SWI indices were within the bounds obtained by Buckingham and Goodale (2010) (the presence of any meaningful effect of group was rejected with all p’s < 0.024, see Supporting Information S1) (Buckingham & Goodale, 2010; Lakens, 2017).

The two CB individuals tested showed SWI indices within the range of the other groups (350g: - 1.145 and -1.034; 700g: -1.507 and -0.646), demonstrating a full strength SWI (Figure 6).

## Methods

### Experiment 2

Experiment 2 was performed as a follow up to the results from Experiment 1, in order to assess the occurrence of the SWI in the absence of haptic size cues. Since we found no group differences in Experiment 1, and our a priori hypothesis was restricted to the effects of transient congenital patterned visual deprivation on the development of the SWI, we ran two groups: CC and SC individuals. Further, given the limitations on recruitment of special populations, as our goal with this experiment was to isolate the visual contribution to the full-sized SWI observed in CC individuals, DC individuals were not tested. In light of the occurrence of a full-sized SWI with (necessarily) exclusively haptic size cues in CB individuals in Experiment 1 and prior studies, we did not repeat an additional haptics-only condition (Ellis and Lederman 1993, Buckingham et al 2015).

Data for this experiment was collected at LVPEI, Hyderabad, India, by the some of the authors, partially with the help of a translator, in English, Hindi or Telugu.

### Participants

The CC group consisted of six individuals defined and classified the same way as in Experiment 1 (referred to as CC: 1 female, 5 males; Age = 17 – 44.7 years, M = 27.67 years, SD = 12.37; Duration of blindness = 2 – 22.01 years, M = 12.98 years, SD = 7.28, Table 1). An additional CC participant was excluded as they did not consistently rate the 350g cubes as less heavy than the 700g cubes, indicating that they were not performing the task in a principled manner, possibly due to translation issues (see Supporting Information S2). For four out of six included participants, visual acuity pre-surgery in the better eye ranged from a minimum of counting fingers at 1m to a maximum of 20/300, with a history of partially absorbed lenses in all four of them. All other criteria, such as presence of nystagmus and family history, pointed towards the presence of dense bilateral cataracts at birth. For the remaining two participants, visual acuity pre-surgery was unknown, but based on the combination of a family history of dense congenital cataracts, very poor visual acuity post-surgery, nystagmus and esotropia, these participants were classified as having dense bilateral congenital cataracts. Visual acuity post-surgery in the better eye in this group ranged from a minimum of 20/400 to a maximum of 20/125.

The SC group consisted of seven individuals with normal or corrected-to-normal vision, with no history of eye injuries or abnormalities (5 females, 2 males; Age = 21-29 years, M = 24.13, SD = 3.08).

### Stimuli and Apparatus

Participants used a white, smooth ribbon to hold and lift one of the same six cubes used in Experiment 1, by pulling it with their dominant hand. The ribbon was wrapped around a metal ring fixed to a wall, in order to minimize friction that could possibly affect weight judgements, and allow participants to estimate the weight of the cube while it was suspended at eye level (Figure 1). An important experimental consideration for the use of a string/handle set-up to test the SWI in sight recovery individuals was to control for the precision of the visual cues provided, due to residual visual impairments in visually impaired individuals (Lewis & Maurer, 2005). Therefore, viewing distance was determined by each CC and SC participant based on how comfortable they were seeing the cube. However, viewing distance was not significantly different between groups (t(4) = 1.656, p = 0.137; CC: Mean = 58 cm, SD = 8.69 cm, Range = 50 – 70 cm; SC: Mean = 64.71 cm, SD = 5.19 cm, Range = 60 – 73 cm). This was done to minimize the possibility of potential differences in illusion size being confounded with differences in visual acuity, due to viewing at a fixed distance. Participants were instructed identically to Experiment 1 described above, and the experimenter placed and removed the cubes from the apparatus for each trial to prevent the participant from having any haptic contact with the stimuli.

Participants were not permitted to haptically handle the cubes at any time, before or during the course of the task, and were naïve to how many sizes and weights were presented. A post-study questionnaire recorded their estimates for how many weights and sizes were presented (see Supporting Information S3).

The same random trial orders used in Experiment 1 were used for Experiment 2, and 60 trials were completed per participant.

### Data Analysis

Z-scores were calculated using a procedure identical to the one described for Experiment 1 above.

The ANOVA model comprised two within-group factors (Size: 3 levels, Weight: 2 levels) and one between groups factor with 2 levels (CC, SC). Post-hoc ANOVAs and t-tests were performed according to the resulting interactions.

Additionally, a cross-experiment ANOVA with group and experimental task as between subject factors and weight as a within subject factor (group: 2 levels for SC and CC; task: 2 levels for Experiment 1 and 2) was performed.

## Results

### 2. Sight recovery individuals show an SWI with exclusively visual size estimates (Experiment 2)

In a group-by-weight-by-size ANOVA performed for Experiment 2, we obtained a main effect of weight (F(1,11) = 110.80, p < 0.001, η_g_ ^2^ = 0.833), indicating that participants performed the task in a principled manner. Additionally, the main effect of size was significant (F(2,22) = 8.82, p = 0.009, η_g_ ^2^ = 0.203), confirming an SWI with this task across groups (Figure 4, Figure 5). The CC participants did not differ from the SC group in their performance on this task, as evidenced by the lack of a significant main effect of group (F(1,11) = 0.158, p = 0.999, η_g_ ^2^ < 0.001), and of significant group-level interactions (group-by-size: F(2,22) = 0.048, p = 0.861, η_g_ ^2^ = 0.002; group-by-weight: F(1,11) = 1.866, p = 0.199, η_g_ ^2^ = 0.077; group-by-size-by-weight: F(2,22) = 0.556, p = 0.509, η_g_ ^2^ = 0.009).

**Figure 4:**
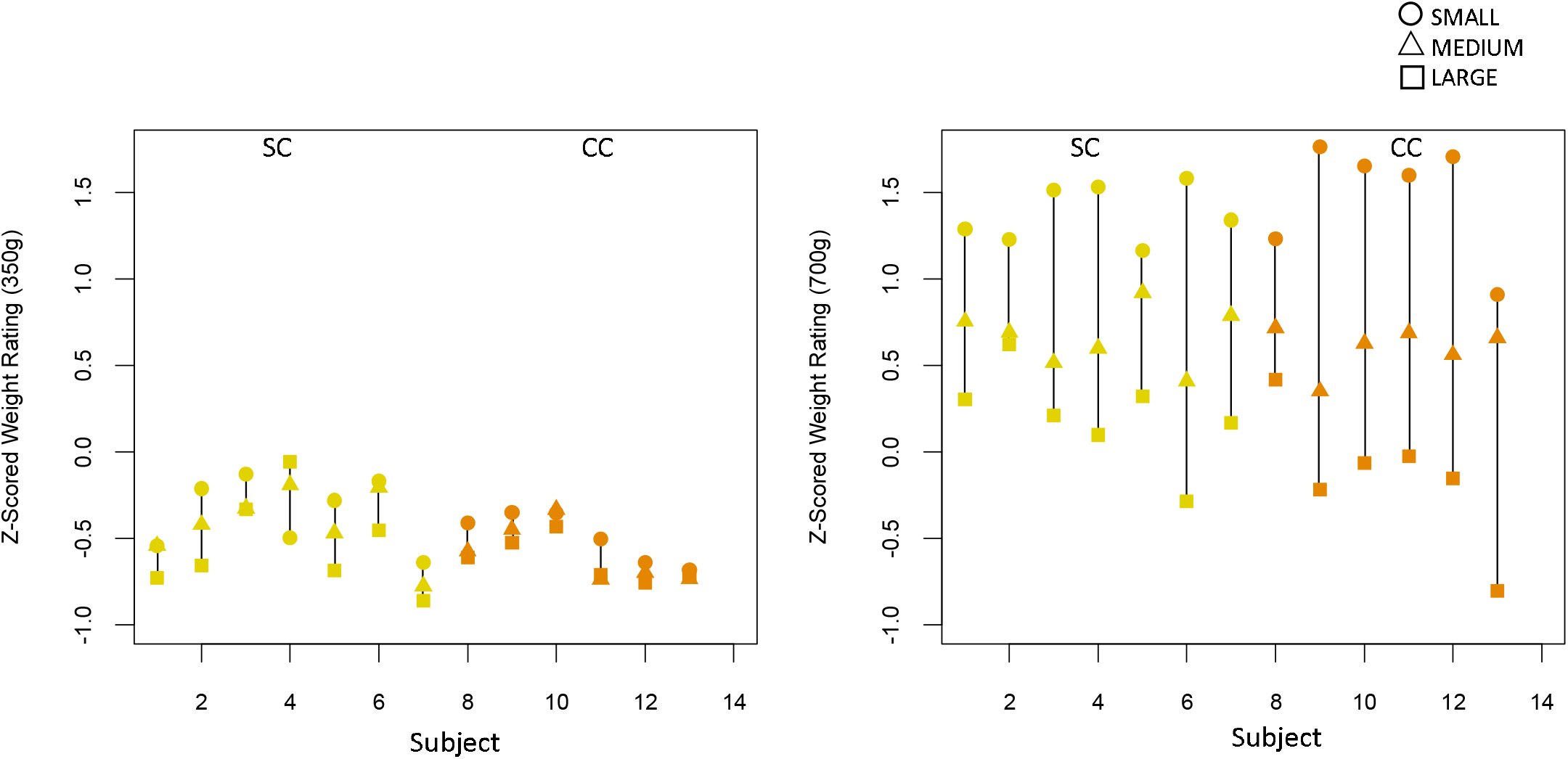
Experiment 2 (with only visual size information); z-scored weight ratings of all individuals in the Sighted Control group (SC, yellow) and Congenital Cataract group (CC, orange) groups (A) for the 350g and (B) for 700g weights (Experiment 2).

**Figure 5:**
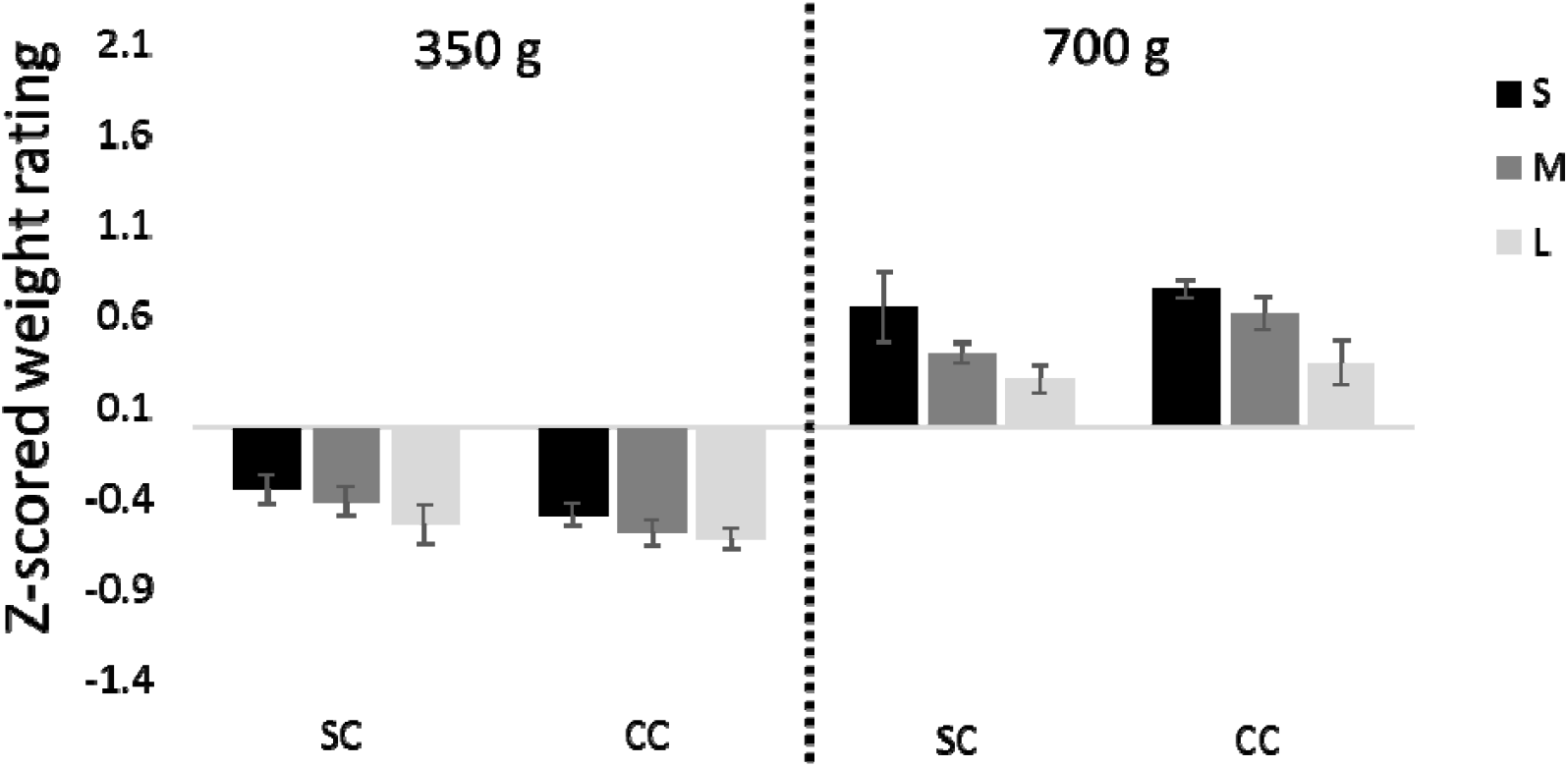
Experiment 2 (with only visual size information); average z-scored weight ratings for the Small (S), Medium (M) and Large (L) cubes for the Sighted Control group (SC) and Congenital Cataract group (CC), for the 350g and 700g weights. Error bars depict standard error of mean (SEM).

Across groups, participants experienced the smaller cube as heavier than the medium cube (350g: t(12) = 2.03, p = 0.056; 700g: t(12) = 2.02, p = 0.066) as well as the large cube (350g: t(12) = 2.74, p = 0.018; 700g: t(12) = 2.86, p = 0.014) of the same weight, and the medium sized cube as heavier than the large cube of the same weight (350g: t(12) = 2.62, p = 0.023; 700g: t(12) = 3.03, p = 0.010), confirming the presence of the SWI with this paradigm.

#### 2.1 SWI Index

The CC and SC groups did not differ in the strength of the illusion for either the 350g (*F*(1,11) = 0.169, *p* = 0.688, η_g_ ^2^ = 0.015), or the 700g weights (*F*(1,11) = 0.001, *p* = 0.971, η_g_ ^2^ < 0.001) (Figure 6). When compared using equivalence testing for the 700g weight, conducted based on available effect sizes in the literature, we found the SWI indices to be equivalent, significantly rejecting any effect of group (both *p*’s < 0.006, see Supporting Information S1).

**Figure 6:**
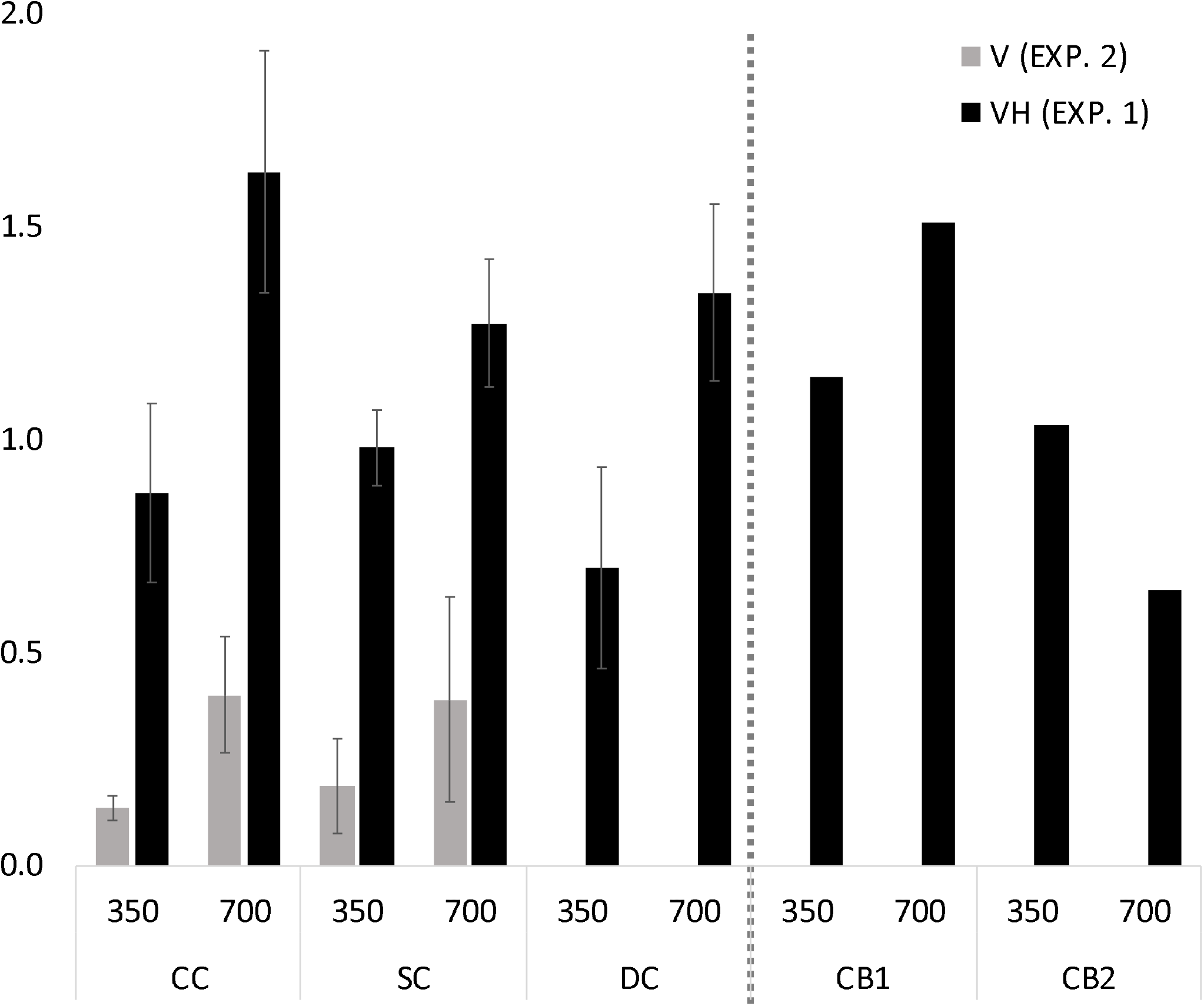
Size-Weight Illusion Indices across groups and for the two CB individuals for Experiment 1 (with visual and haptic size information, black bars) and Experiment 2 (with only visual size information, gray bars), for the 350g and 700g weights. The y-Axis depicts the difference in z-scores between the large and small cubes of each weight, averaged for each group (CC, DC, and SC). Error bars depict the SEM.

### 3. The visual contribution to the SWI is identical in sight recovery and sighted individuals

We additionally compared the SC and CC groups of Experiment 1 and 2 in a group-by-weight-by-experimental task ANOVA, in order to assess whether the groups differed between how they performed when both visual and haptic size estimates were available, compared to when only visual size information was available. We confirmed that the SWI was significantly stronger in Experiment 1 compared to Experiment 2 (main effect of Experimental Task: *F*(1,26) = 56.396, *p* < 0.001, η_g_ ^2^ = 0.511), and stronger for the 700g than the 350g weight (main effect of Weight: *F*(1,26) = 9.089, *p* = 0.006, η_g_ ^2^ = 0.153). The group-by-experiment interaction (*F*(1,26) = 0.357, *p* = 0.555, η_g_ ^2^ = 0.006), group-by-weight interaction (*F*(1,26) = 1.083, *p* = 0.307, η_g_ ^2^ = 0.021), and group-by-weight-by-experiment interaction were all non-significant (*F*(1,26) = 0.631, *p* = 0.424, η_g_ ^2^ = 0.012), confirming that in both experiments CC individuals perceived an indistinguishably strong SWI from SC individuals, suggesting that the “visual”contribution to the SWI did not differ in the two groups.

#### 3.1 Relationship between strength of the SWI and the duration of visual deprivation

To test for a possible effect of duration of patterned visual deprivation on the strength of the SWI, we calculated the correlation between age at surgery and the average SWI Index (across 350g and 700g) for CC individuals. This correlation was not significant neither for Experiment 1 (r=0.05, *t*(4)=0.10, *p*=0.463), nor did the correlation reach significance for Experiment 2 (r = -0.76, *t*(4) = -2.357, *p* = 0.078). Additionally, no correlation was observed between illusion size and age at time tested in either group (see Supporting Information S4.3).

## Discussion

The present study investigated whether the manifestation of the size-weight illusion (SWI) depends on patterned visual experience after birth. We tested sight recovery individuals with a history of dense bilateral congenital cataracts, and compared this group to sight recovery individuals with a history of developmental cataracts, as well as a group of normally sighted controls. Our results demonstrated a significant “classical”SWI (with visual and haptic size information available) in all groups; indeed, the size of the SWI was indistinguishable between the three groups. Furthermore, we replicated previous results from Ellis and Lederman and showed that two permanently congenitally blind individuals experienced the SWI to a degree that fell within the range of all the other participants (Ellis & Lederman, 1993).

We additionally used a string set-up in Experiment 2 to test whether the CC group used visual size cues, rather than relying only on haptic size cues in Experiment 1 (Buckingham et al., 2015). As in Experiment 1, the SWI experienced with exclusively visual size information was equivalent across sight recovery individuals with a history of congenital cataracts, and sighted controls. Together, these results suggest that the visuo-haptic SWI is resilient to atypical visual experience after birth.

### No sensitive period effects for visuo-haptic integration as tested by the SWI

A previous study of individuals who were operated upon for dense bilateral congenital cataracts reported that in an object matching task conducted two days-post-surgery - while unimodal tactile and visual performance was observed to be at ceiling, tactile to visual mapping was found to be severely impaired (Held et al., 2011, n = 5). However, the authors observed that this ability rapidly improved over the next five days. A subsequent case study of sight recovery suggested that visuo-tactile processing recovers in object recognition and object matching tasks within three days of sight restoration, despite a lack of visual experience after birth (Chen et al., 2016, n = 1). As these studies tested participants closer to the date of surgery, and given that the sight recovery individuals in the present study were all tested one year from surgery in order to exclude acute but transient surgery effects, our findings are consistent with these existing studies on visuo-haptic object recognition. However, both Held et al and Chen et al tested visuo-haptic transfer through object matching tasks. By contrast, we provide evidence for the recovery of visuo-haptic *integration* (i.e. a unified percept by fusing input from both sensory modalities) despite early patterned visual deprivation, therefore extending these studies (Singh et al., 2018; Stein et al., 2010). Our findings might be considered surprising in light of two prospective studies, which suggested a protracted developmental pathway for the SWI. I. The first study observed that the SWI increased in size after the age of 5 years (Chouinard et al., 2019) They related this increase to the development of abstract reasoning skills. However, abstract reasoning explained no more than about 10% of the effect, and the SWI existed even in the youngest group. A second study showed that typically developing children did not optimally integrate visuo-haptic input in an adult-like manner until the age of 10 years (Gori et al., 2008). However, optimal integration is typically defined as optimal cue integration as predicted by forced fusion models. These models weight individual cues according to their relative reliability to derive a multisensory outcome. It has more recently been demonstrated that in situations where it is ambiguous whether or not to integrate sensory information, the data from children as young as 5 years of age, like those of adults, are better explained by causal inference models (Rohlf, Li, Bruns, & Röder, 2020). Given that some (n=4) of our CC participants had been older than 10 years of age at the time of surgery, our results suggest that patterned vision during this period of multisensory development was not crucial for the typical manifestation of the SWI with either visuo-haptic or only visual size information.

These results showing an indistinguishable SWI in sight recovery individuals, both with a congenital as well as a developmental history of transient blindness, provide evidence that the multisensory processes underlying the SWI are resilient to atypical visual experience. First, participants of both cataract groups still suffered visual impairments at the time of testing. Nevertheless, the SWI was not smaller in magnitude in either group, compared to normally sighted individuals. A smaller SWI in sight recovery individuals would have been expected from an the aforementioned reliability-based optimal integration account (Ernst & Banks, 2002). Second, neither years of blindness nor the timing of the transient phase of blindness (developmental vs. congenital) had a significant influence on the size of the SWI. Finally, the extent of the “visual”contribution to the SWI was indistinguishable between sight recovery and sighted individuals. This identical behavioral performance, regardless of atypical visual history across groups and tasks, provides strong evidence that visuo-haptic processing, as assessed with the SWI, does not rely on sensitive period plasticity to develop normally.

It is possible that different underlying neural mechanisms support the identically sized SWI in sight recovery individuals (Bedny, 2017). As sensitive periods are properties of neural circuits, further neuroimaging studies need to confirm whether visuo-haptic processing develops normally in the absence of typical visual experience (Knudsen, 2004; Takesian & Hensch, 2013). Additionally, the absence of sensitive period effects in one tested behavior does not contradict the general role of sensitive period plasticity (Hadad, Maurer, & Lewis, 2012; Lewis & Maurer, 2005; Röder, Kekunnaya, & Guerreiro, 2020; Röder, Ley, Shenoy, Kekunnaya, & Bottari, 2013; Takesian & Hensch, 2013). Disengaging functions which do and do not develop within sensitive periods will, in the long run, be essential to uncovering the general principles of functional brain development.

To the best of our knowledge, the present study is the first to conclusively show the manifestation of a full-strength SWI in sight recovery individuals, both for the condition with visual and haptic size information, as well as the condition with only visual size information. Further, strict criteria were used for the inclusion of sight recovery participants, in order to ensure high homogeneity of etiology within the CC and DC group. We chose a retrospective, developmental approach with individuals who underwent cataract reversal surgery before the age of 23 years, and were older than 8 years of age at the time of testing. After the age of 8 years, no further increase in the SWI had been observed in prospective studies (Chouinard et al., 2019). The present study did not find any difference in the size of the SWI, neither for the visuo-haptic nor for the visual experiment. A significant, equivalent SWI was consistently perceived by individual participants across all groups, despite the fact that in the cataract groups, age at surgery, time since surgery and visual acuity at time of testing varied, potentially increasing between group differences. Our stringent inclusion criteria restricted the sample size within a special population, however, all individual subjects showed the SWI with established paradigms in a consistent pattern (Figures 2,3), allowing us to interpret the lack of group differences and confirming them with equivalence testing. We consider these results to be highly robust evidence against the dependence of the SWI on early visual input, therefore suggesting the lack of a sensitive period for the development of the SWI.

### Mechanisms of the SWI

What are the possible mechanisms by which the SWI could manifest in CC individuals? While our study does not allow us to disentangle the models explaining the occurrence of the SWI, interpreting our results in light of these hypotheses can shed light on the mechanisms of the SWI. Below, we engage with the dominant models of the SWI.

On one hand, it could be assumed that the SWI is a purely haptic process, as was consistent with earlier reports of the SWI manifesting in congenitally blind adults, and replicated in two congenitally bind adults in the present study (Ellis & Lederman, 1993; Rice, 1898). However, a purely haptic account of the SWI would have predicted the absence of the SWI when haptic size cues are unavailable to sight recovery individuals. In fact, prior studies employing a string set-up in congenitally blind individuals, as expected, did not observe an SWI (Buckingham et al., 2015; Ellis & Lederman, 1993). Instead, in the present study, sight recovery individuals perceived an SWI even when only visual size cues were available. Additionally, a recent study observed that congenitally blind individuals experience the SWI without haptic size cues, but when size estimates were obtained through echolocation (Buckingham et al., 2015). Together, this evidence strongly argues against an exclusively haptic account of the SWI.

On the other hand, it has been suggested that the SWI is a multisensory phenomenon occurring due to a conflict between concurrent visual (size) and haptic (weight) sensory input (Dijker, 2014; Grandy & Westwood, 2006; Kawai et al., 2007). Within this framework, our data suggest that while the SWI can develop through haptic input alone, it might nevertheless be modulated by visual input, due to the recovery of visuo-haptic processing despite atypical visual experience. (Chen et al., 2016; Chen et al., 2017; Held et al., 2011) Indeed, full or partial recovery of visuo-tactile functions have been reported, depending on the task. While prior studies have shown that in a simultaneity judgement task designed to test a unified multisensory percept, visuo-tactile performance was unimpaired despite a lack of early visual experience (Chen et al., 2017; Putzar et al., 2012), sight recovery individuals did not show normal visuo-tactile temporal order biases (Badde et al., 2020; Ley et al., 2013). Additionally, our findings of a larger SWI when both visual and haptic size information was available than when only a visual size estimate was possible, in both sight recovery and sighted individuals, fit with a multisensory framework for the occurrence of the SWI (Ellis & Lederman, 1993; Pick & Pick, 1967).

## Conclusions

The occurrence of the Size-Weight Illusion (SWI), both when visual and haptic size information was available, as well as when only visual size information was assessable, was resilient to atypical visual experience within the first months and years of life. These results provide strong evidence that the visuo-haptic processes underlying the SWI do not require typical visual experience within a sensitive period for normal development. Further studies are needed to explore whether the SWI is supported by the same neural mechanisms in typical and atypical development by employing neuroscience techniques (Chouinard et al., 2009).

### Data Availability Statement

The datasets generated/analyzed during this study are available from the corresponding author upon reasonable request.

## Competing Interests Statement

The authors declare no competing interests.

## Author Contributions

RP designed and collected data for Experiment 2, analyzed the data for both experiments, made the figures and tables and wrote the paper. MG, PL and DB designed and collected data for Experiment 1, MG analyzed the data for both experiments and wrote the paper. IS recruited, counselled and diagnosed sight recovery individuals and assisted in data collection for Experiment 2. RK counselled and diagnosed sight recovery individuals for all experiments and supervised the work. BR designed the study, collected the data for Experiment 1 and wrote the paper. All authors edited the manuscript.

## Acknowledgements

We are grateful to D. Balasubramanian and the LVPEI for supporting the study at the LVPEI in Hyderabad. We thank Kabilan Pitchaimuthu, Sven Leach and Suddha Sourav for helping with the data acquisition. Dirk Waschatz provided technical assistance. The study was funded by the German Research Foundation (DFG Ro 2625/10-1) and the European Research Council grant ERC-2009-AdG 249425-CriticalBrainChanges to Brigitte Röder. Rashi Pant was supported by a PhD student fellowship from the Hector Fellow Academy gGmbH.

## Notes

### Competing Interest Statement

The authors have declared no competing interest.

## References

Azañón, E., Camacho, K., Morales, M., & Longo, M. R. (2018). The Sensitive Period for Tactile Remapping Does Not Include Early Infancy. Child Development. https://doi.org/10.1111/cdev.12813

Badde, S., Ley, P., Rajendran, S. S., Shareef, I., Kekunnaya, R., & Röder, B. (2020). Sensory experience during early sensitive periods shapes cross-modal temporal biases. ELife. https://doi.org/10.7554/ELIFE.61238

Bedny, M. (2017). Evidence from Blindness for a Cognitively Pluripotent Cortex. Trends in Cognitive Sciences. https://doi.org/10.1016/j.tics.2017.06.003

Birch, E. E., Stager, D., Leffler, J., & Weakley, D. (1998). Early treatment of congenital unilateral cataract minimizes unequal competition. Investigative Ophthalmology and Visual Science.

Buckingham, G., & Goodale, M. A. (2010). Lifting without seeing: The role of vision in perceiving and acting upon the size weight Illusion. PLoS ONE. https://doi.org/10.1371/journal.pone.0009709

Buckingham, G., Milne, J. L., Byrne, C. M., & Goodale, M. A. (2015). The Size-Weight Illusion Induced Through Human Echolocation. Psychological Science. https://doi.org/10.1177/0956797614561267

Chen, J., Wu E. De Chen, X., Zhu, L. H., Li, X., Thorn, F., … Qu, J. (2016). Rapid Integration of Tactile and Visual Information by a Newly Sighted Child. Current Biology. https://doi.org/10.1016/j.cub.2016.02.065

Chen, Y. C., Lewis, T. L., Shore, D. I., & Maurer, D. (2017). Early Binocular Input Is Critical for Development of Audiovisual but Not Visuotactile Simultaneity Perception. Current Biology. https://doi.org/10.1016/j.cub.2017.01.009

Chouinard, P. a., Large, M. E., Chang, E. C., & Goodale, M. a. (2009). Dissociable neural mechanisms for determining the perceived heaviness of objects and the predicted weight of objects during lifting: An fMRI investigation of the size-weight illusion. NeuroImage. https://doi.org/10.1016/j.neuroimage.2008.08.023

Chouinard, P. A., Matheson, K. G., Royals, K. A., Landry, O., Buckingham, G., Saccone, E. J., & Hocking, D. R. (2019). The development of the size–weight illusion in children coincides with the development of nonverbal cognition rather than motor skills. Journal of Experimental Child Psychology. https://doi.org/10.1016/j.jecp.2019.03.006

de Heering, A., Dormal, G., Pelland, M., Lewis, T., Maurer, D., & Collignon, O. (2016). A Brief Period of Postnatal Visual Deprivation Alters the Balance between Auditory and Visual Attention. Current Biology. https://doi.org/10.1016/j.cub.2016.10.014

Dijker, A. J. M. (2014). The role of expectancies in the size-weight illusion: A review of theoretical and empirical arguments and a new explanation. Psychonomic Bulletin and Review. https://doi.org/10.3758/s13423-014-0634-1

Eagleman, D. M. (2001). Visual illusions and neurobiology. Nature Reviews Neuroscience. https://doi.org/10.1038/35104092

Ellis, R. R., & Lederman, S. J. (1993). The role of haptic versus visual volume cues in the size-weight illusion. Perception & Psychophysics. https://doi.org/10.3758/BF03205186

Ernst, M. O., & Banks, M. S. (2002). Humans integrate visual and haptic information in a statistically optimal fashion. Nature. https://doi.org/10.1038/415429a

Flanagan, J. R., Bittner, J. P., & Johansson, R. S. (2008). Experience Can Change Distinct Size-Weight Priors Engaged in Lifting Objects and Judging their Weights. Current Biology. https://doi.org/10.1016/j.cub.2008.09.042

Furth, H. G. (1961). Effect of Training on the Adaptation Level of the Size-Weight Illusion with Normal, Deaf, and Blind Subjects. Perceptual and Motor Skills, 13, 155–160. https://doi.org/10.2466/pms.1961.13.2.155

Gandhi, T., Kalia, A., Ganesh, S., & Sinha, P. (2015). Immediate susceptibility to visual illusions after sight onset. Current Biology. https://doi.org/10.1016/j.cub.2015.03.005

Gori, M., Del Viva, M., Sandini, G., & Burr, D. C. (2008). Young Children Do Not Integrate Visual and Haptic Form Information. Current Biology. https://doi.org/10.1016/j.cub.2008.04.036

Grandy, M. S., & Westwood, D. A. (2006). Opposite perceptual and sensorimotor responses to a size-weight illusion. Journal of Neurophysiology. https://doi.org/10.1152/jn.00851.2005

Guerreiro, M. J. S., Putzar, L., & Roder, B. (2016). The Effect of Early Visual Deprivation on the Neural Bases of Auditory Processing. Journal of Neuroscience, 36(5), 1620–1630. https://doi.org/10.1523/JNEUROSCI.2559-15.2016

Guerreiro, Maria J.S., Putzar, L., & Röder, B. (2015). The effect of early visual deprivation on the neural bases of multisensory processing. Brain. https://doi.org/10.1093/brain/awv076

Guerreiro, Maria J.S., Putzar, L., & Röder, B. (2016). Persisting Cross-Modal Changes in Sight-Recovery Individuals Modulate Visual Perception. Current Biology. https://doi.org/10.1016/j.cub.2016.08.069

Hadad, B. S., Maurer, D., & Lewis, T. L. (2010). The development of contour interpolation: Evidence from subjective contours. Journal of Experimental Child Psychology. https://doi.org/10.1016/j.jecp.2010.02.003

Hadad, B. S., Maurer, D., & Lewis, T. L. (2012). Sparing of sensitivity to biological motion but not of global motion after early visual deprivation. Developmental Science. https://doi.org/10.1111/j.1467-7687.2012.01145.x

Hadad, B. S., Maurer, D., & Lewis, T. L. (2017). The role of early visual input in the development of contour interpolation: the case of subjective contours. Developmental Science. https://doi.org/10.1111/desc.12379

Held, R., Ostrovsky, Y., Degelder, B., Gandhi, T., Ganesh, S., Mathur, U., & Sinha, P. (2011). The newly sighted fail to match seen with felt. Nature Neuroscience. https://doi.org/10.1038/nn.2795

Kawai, S., Henigman, F., MacKenzie, C. L., Kuang, A. B., & Faust, P. H. (2007). A reexamination of the size-weight illusion induced by visual size cues. Experimental Brain Research. https://doi.org/10.1007/s00221-006-0803-1

Knudsen, E. I. (2004). Sensitive periods in the development of the brain and behavior. J Cogn Neurosci, 16(8), 1412–1425. https://doi.org/10.1162/0898929042304796

Kumar, G. V., Halder, T., Jaiswal, A. K., Mukherjee, A., Roy, D., & Banerjee, A. (2016). Large scale functional brain networks underlying temporal integration of audio-visual speech perception: An eeg study. Frontiers in Psychology. https://doi.org/10.3389/fpsyg.2016.01558

Lakens, D. (2017). Equivalence Tests: A Practical Primer for t Tests, Correlations, and Meta-Analyses. Social Psychological and Personality Science. https://doi.org/10.1177/1948550617697177

Lewis, T. L., & Maurer, D. (2005). Multiple sensitive periods in human visual development: Evidence from visually deprived children. Developmental Psychobiology, 46(3), 163–183. https://doi.org/10.1002/dev.20055

Lewkowicz, D. J., & Röder, B. (2015). The effects of experience on the development of multisensory processing. The New Handbook of Multisensory Processes, (September).

Ley, P., Bottari, D., Shenoy, B. H., Kekunnaya, R., & Röder, B. (2013). Partial recovery of visual-spatial remapping of touch after restoring vision in a congenitally blind man. Neuropsychologia. https://doi.org/10.1016/j.neuropsychologia.2013.03.004

Masin, S. C., & Crestoni, L. (1988). Experimental demonstration of the sensory basis of the size-weight illusion. Perception & Psychophysics. https://doi.org/10.3758/BF03210411

Maurer, D. (2017). Critical periods re-examined: Evidence from children treated for dense cataracts. Cognitive Development. https://doi.org/10.1016/j.cogdev.2017.02.006

McKyton, A., Ben-Zion, I., Doron, R., & Zohary, E. (2015). The Limits of Shape Recognition following Late Emergence from Blindness. Current Biology. https://doi.org/10.1016/j.cub.2015.06.040

McKyton, A., Ben-Zion, I., & Zohary, E. (2018). Lack of Automatic Imitation in Newly Sighted Individuals. Psychological Science. https://doi.org/10.1177/0956797617731755

Meredith, M., & Stein, B. (1983). Interactions among converging sensory inputs in the superior colliculus. Science. https://doi.org/10.1126/science.6867718

Murray, D. J., Ellis, R. R., Bandomir, C. A., & Ross, H. E. (1999). Charpentier (1891) on the size-weight illusion. Perception & Psychophysics. https://doi.org/10.3758/BF03213127

Nath, A. R., & Beauchamp, M. S. (2012). A neural basis for interindividual differences in the McGurk effect, a multisensory speech illusion. NeuroImage. https://doi.org/10.1016/j.neuroimage.2011.07.024

Peters, M. A. K., Balzer, J., & Shams, L. (2015). Smaller = denser, and the brain knows it: Natural statistics of object density shape weight expectations. PLoS ONE. https://doi.org/10.1371/journal.pone.0119794

Pick, H. L., & Pick, A. D. (1967). A developmental and analytic study of the size-weight illusion. Journal of Experimental Child Psychology. https://doi.org/10.1016/0022-0965(67)90064-1

Plaisier, M. A., & Smeets, J. B. J. (2015). Object size can influence perceived weight independent of visual estimates of the volume of material. Scientific Reports. https://doi.org/10.1038/srep17719

Putzar, L., Goerendt, I., Heed, T., Richard, G., Büchel, C., & Röder, B. (2010). The neural basis of lip-reading capabilities is altered by early visual deprivation. Neuropsychologia. https://doi.org/10.1016/j.neuropsychologia.2010.04.007

Putzar, L., Goerendt, I., Lange, K., Rösler, F., & Röder, B. (2007). Early visual deprivation impairs multisensory interactions in humans. Nature Neuroscience. https://doi.org/10.1038/nn1978

Putzar, L., Gondan, M., & Rder, B. (2012). Basic multisensory functions can be acquired after congenital visual pattern deprivation in humans. Developmental Neuropsychology. https://doi.org/10.1080/87565641.2012.696756

Putzar, L., Hötting, K., & Röder, B. (2010). Early visual deprivation affects the development of face recognition and of audio-visual speech perception. Restorative Neurology and Neuroscience. https://doi.org/10.3233/RNN-2010-0526

Putzar, L., Hötting, K., Rösler, F., & Röder, B. (2007). The development of visual feature binding processes after visual deprivation in early infancy. Vision Research. https://doi.org/10.1016/j.visres.2007.07.002

Rice, J. (1898). The size–weight illusion among the blind. Amm Psychol Assn, (27), 81–87.

Röder, B., Kekunnaya, R., & Guerreiro, M. J. S. (2020). Neural mechanisms of visual sensitive periods in humans. Neuroscience & Biobehavioral Reviews. https://doi.org/10.1016/j.neubiorev.2020.10.030

Röder, B., Ley, P., Shenoy, B. H., Kekunnaya, R., & Bottari, D. (2013). Sensitive periods for the functional specialization of the neural system for human face processing. Proceedings of the National Academy of Sciences of the United States of America. https://doi.org/10.1073/pnas.1309963110

Rohlf, S., Li, L., Bruns, P., & Röder, B. (2020). Multisensory Integration Develops Prior to Crossmodal Recalibration. Current Biology. https://doi.org/10.1016/j.cub.2020.02.048

Ross, H. E. (1966). Sensory information necessary for the size-weight illusion [58]. Nature. https://doi.org/10.1038/212650a0

Sams, M., Aulanko, R., Hamalainen, M., Hari, R., Lounasmaa, O. V, Lu, S.-T., & Simola, J. (1991). Seeing speech: Visual information in the human auditory cortex. Neuroscience Letters.

Singh, A. K., Phillips, F., Merabet, L. B., & Sinha, P. (2018). Why Does the Cortex Reorganize after Sensory Loss? Trends in Cognitive Sciences. https://doi.org/10.1016/j.tics.2018.04.004

Stein, B. E., Burr, D., Constantinidis, C., Laurienti, P. J., Alex Meredith, M., Perrault, T. J., … Lewkowicz, D. J. (2010). Semantic confusion regarding the development of multisensory integration: A practical solution. European Journal of Neuroscience. https://doi.org/10.1111/j.1460-9568.2010.07206.x

Takesian, A. E., & Hensch, T. K. (2013). Balancing plasticity/stability across brain development. In Progress in Brain Research. https://doi.org/10.1016/B978-0-444-63327-9.00001-1

World Health Organisation. (2019). World report on vision. World health Organisation.

