## Supporting Information for "The Size-Weight Illusion is unimpaired in individuals with a history of congenital visual deprivation"

#### S1. Equivalence Testing

##### S1.1 Experiment 1.

In order to clarify the non-significant group effects (i.e. the SWI being equally strong in individuals with atypical visual experience as in normally sighted controls), we used post-hoc equivalence testing to compare the SWI indices between the CC, DC and SC groups (Lakens, 2017; Lakens, Scheel, & Isager, 2018). For this purpose, we chose to define the upper and lower bounds of interest based on published values obtained by Buckingham and Goodale (2010), i.e. the study we had used as a model for the present experimental procedure. They had tested a relatively large sample size of healthy individuals (n=39), calculated the SWI index ("effect size") similar to our method, and reported z-scored values in their Figure 1 (Buckingham & Goodale, 2010). In an additional literature search, we were not able to detect any additional study analyzing and reporting group differences in the SWI, as assessed via a comparable paradigm to the one we had implemented. Thus, effect sizes for group differences were not available in the literature. Further, we report only the values we obtained for the 700g weight as the selected paper used an identical weight (exploratory equivalence testing with the 350g condition produced, however, identical results). We used WebPlotDigitizer (v 3.9) in order to select conservative values (Rohatgi, 2015; Rohatgi, 2010). The lowest possible value for the small cube and the highest possible value for the large cube were selected for the 700g weight condition, which reflects the smallest possible effect size. We used these values to calculate an approximate mean SWI index (M = -1.782, SD = 1.134), and an upper and lower bound corresponding to a 98% confidence interval (CI) of this normal distribution (0.859, -4.424).

Equivalence testing was performed using the TOSTER package (https://github.com/Lakens/TOSTER) in R (Version: 3.3.2). We compared the average SWI index between the SC and CC, SC and DC and CC and DC groups to determine the presence of an effect outside of the bounds obtained, for the 700g weight (Table S1). Equivalence testing assumes the presence of such an effect between the compared means of the compared groups, and therefore equivalence is defined as the significant rejection of the null hypothesis (Lakens, 2017). We found that the SWI effect measured by the SWI index was equivalent between all groups compared – i.e. the presence of an effect of group that lies outside the described upper or lower bounds was rejected with $p$'s < 0.024 (Table S1).

##### S1.2 Experiment 2.

Using an identical procedure to Experiment 1, when we compared CC and SC individuals on their illusion strength for the 700g weight, the effect size was found to be equivalent (Lower Bound: $t(11) = 2.99$, $p = 0.006$; Upper Bound: $t(11) = -15.22$, $p < 0.001$, Table S1).

**S2. Excluded participant (Experiment 2)**

One CC participant (36, Female) was excluded from data analysis on account of not cooperating, and the experiment was aborted after 30 trials. During the experiment, it was observed that she was seemingly randomly giving objects one of four possible numerical weights, even when it was reiterated that she should try to focus on the perceived weight of the object and could freely use any scale. She failed to rate the 700g medium cube as heavier than the 350g medium cube, confirming that she was not assigning principled weight ratings. Based on this, her data was excluded from the final analysis.

**S3. Post-Study Questionnaire (Experiment 2)**

After the experiment was complete, we asked participants how many distinct weights and sizes they perceived across the 60 trials.

The estimate for the number of cube sizes presented ranged from 3-10 (Mean = 5.8, Median = 3 ) in the CC and 3-7 (Mean = 4.29, Median = 4) in the SC group. Participants of both groups were subjectively asked to answer how well they could see during the task, and all participants across both groups stated that they could see the stimuli adequately well.

The estimate for the number of weights presented ranged from 5-20 (Mean = 12.33, Median = 9.5) in the CC group and 5-8 (Mean = 6.28, Median = 6) in the SC group, confirming that none of the participants perceived only two weights during the task, and demonstrating the robustness of the SWI perceived by both groups with this paradigm.

**S4. Correlations**

**S4.1 Duration of Blindness (Experiments 1 and 2).**

For Experiment 2, to disambiguate the trending correlation between duration of visual deprivation and SWI Index in CC individuals, we further tested a multiple regression with duration of visual deprivation as well as visual acuity as predictor variables (no autocorrelation between variables, $r = -0.125$, $t(4) = -0.253$, $p = 0.813$). No significant effect of duration of deprivation ($t(4) = -2.326$, $p = -0.102$)

or visual acuity ($t(4) = 1.224$, $p = 0.308$) was observed, indicating that the variance in the SWI Index was not significantly explained by duration of visual deprivation (Figure S1).

**S4.2 Pre-surgery visual acuity (Experiment 2)**

In Experiment 2, to ensure that better vision as a result of absorbed lenses prior to surgery did not have an impact on the SWI, we tested for a relationship between SWI Index and pre-surgery visual acuity. Of the 4 participants with a known pre-surgery visual acuity, 3 participants had a quantifiable value (i.e. better than CF), based on which there was no significant correlation between pre-surgery acuity and SWI size ($r = -0.580$, $t = -0.711$, $p = 0.303$).

**S4.3 Age (Experiment 2)**

To ensure that we did not miss potential differences in SWI size due to age differences between our groups, we tested for any effect of age on SWI size in our sample. We observed no significant correlation between age and SWI index in either group (Figure S2).

**S4.4 Time since surgery (Experiments 1 and 2)**

Additionally, we also tested for any effect of time since surgery on SWI size in our sample. This was calculated by subtracting the age at first surgery from the age on date tested, for every CC participant, and correlated with the mean SWI index. We observed no significant correlation between time since surgery and SWI index in Experiment 1 or 2 (Figure S3).

### References

Buckingham, G., & Goodale, M. A. (2010). Lifting without seeing: The role of vision in perceiving and acting upon the size weight Illusion. *PLoS ONE*. https://doi.org/10.1371/journal.pone.0009709

Lakens, D. (2017). Equivalence Tests: A Practical Primer for t Tests, Correlations, and Meta-Analyses. *Social Psychological and Personality Science*. https://doi.org/10.1177/1948550617697177

Lakens, D., Scheel, A. M., & Isager, P. M. (2018). Equivalence Testing for Psychological Research: A Tutorial. *Advances in Methods and Practices in Psychological Science*. https://doi.org/10.1177/2515245918770963

Rohatgi, A. (2010). WebPlotDigitizer - extract data from plots, images, and maps. *Arohatgi*.

Rohatgi. (2015). WebPlotDigitizer 3.9. Retrieved from http://arohatgi.info/WebPlotDigitizer

### Supplementary Figures and Tables

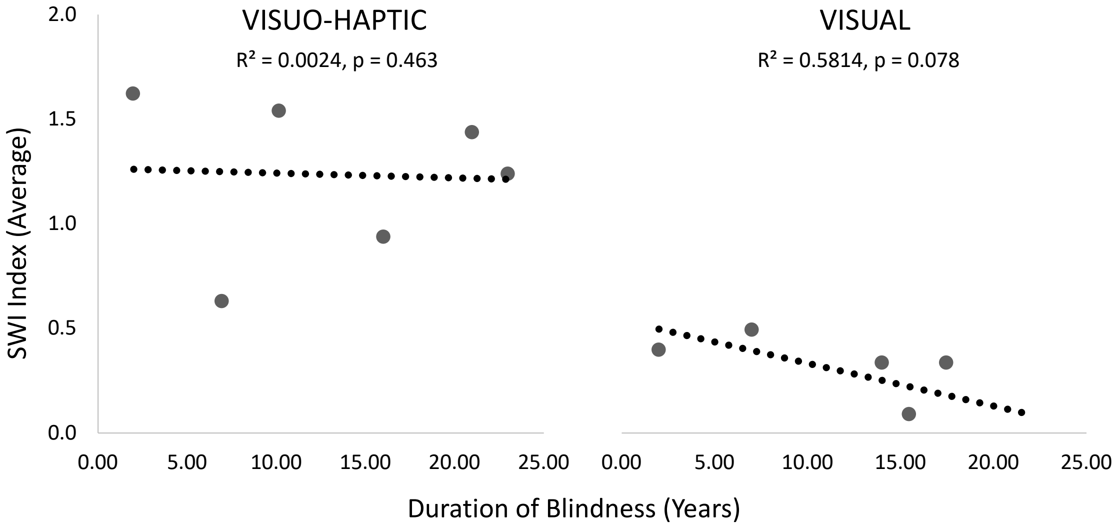


*Fig S1: Correlation between duration of blindness at the time of testing and the calculated SWI Index for the CC group in Experiment 1 (left) and Experiment 2 (right).*

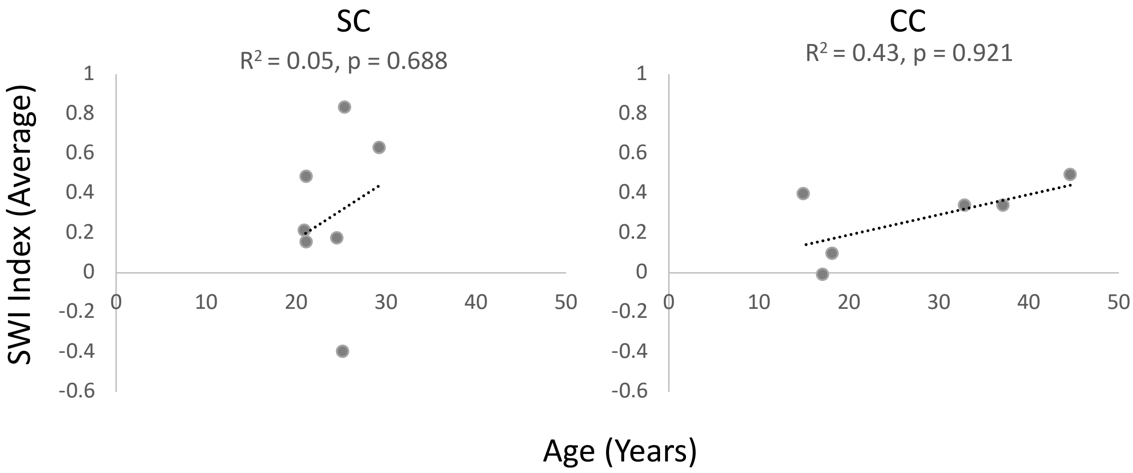


*Figure S2: Correlation between age in years and SWI Index for Experiment 2.*

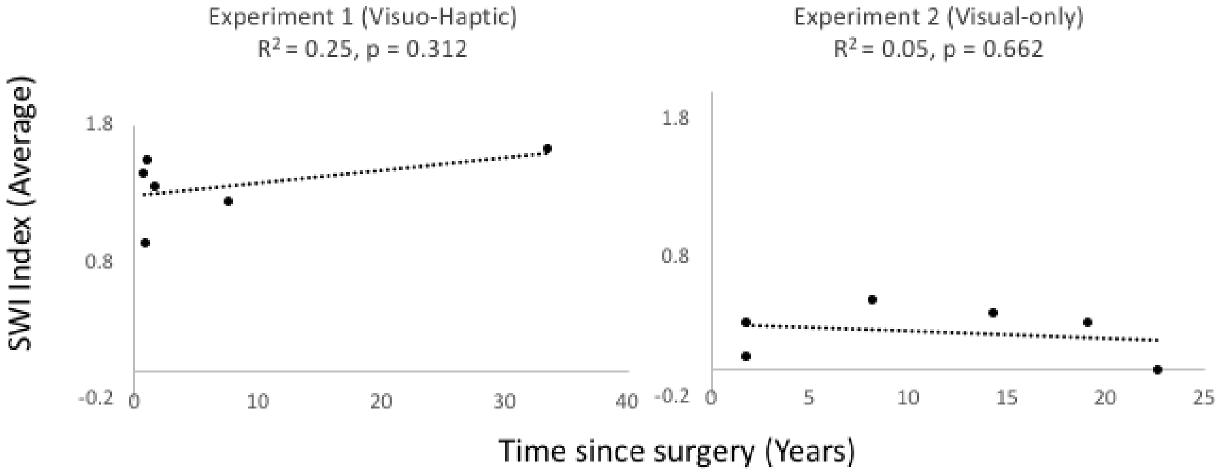


*Figure S3: Correlation between time since surgery in years and SWI Index for Experiments 1 and 2.*

| **Bound** | **CC \| SC** | | **DC \| SC** | | **CC \| DC** | |
|---|---|---|---|---|---|---|
| | ***t(15)*** | ***p*** | ***t(17)*** | ***p*** | ***t(14)*** | ***p*** |
| **Lower** | 13.74 | < 0.001 | 17.16 | < 0.001 | 14.79 | <0.001 |
| **Upper** | -4.09 | < 0.001 | -3.66 | 0.001 | -2.19 | 0.023 |

*Table S1: Equivalence test results comparing mean SWI Indices (effect sizes) between the SC, the CC and the DC groups for the 700g weight. Results of equivalence testing determine the likelihood of rejecting the null hypothesis (i.e. the presence of an effect size difference between the tested groups, outside of the specified bounds). T and p values reported are for the one-sided t-test against each of the two bounds.*
